# A real-time detection and non-destructive warning method for zebrafish body surface anomalies based on improved YOLOv8 framework

**DOI:** 10.1101/2025.04.28.649888

**Authors:** Danying Cao, Hong Yang, Cheng Guo, Yingyin Cheng, Wanting Zhang, Mijuan Shi, Xiao-Qin Xia

## Abstract

The detection of anomalies on fish surfaces is of critical importance for assessing fish health status, preventing fish disease outbreaks, predicting changes in water quality, enhancing fish welfare. The zebrafish (*Danio rerio*), a key model organism, has been increasingly utilized in various fields, including medicine, genetics and environmental toxicology. This has led to a corresponding increase in demand for intelligent management and detection systems. However, traditional methods of fish disease detection may have irreversible effects on fish, particularly small species, and often fail to meet the precision, non-destructive warning, and real-time requirements for zebrafish detection. To address this issue, this study proposes a novel method based on the YOLOv8 framework, designated ESC-YOLOv8-seg. This method significantly enhances the precision and speed of detecting surface abnormalities on small fish in complex settings by integrating the EMA the SPPELAN and C2f-Faster modules, and adding a detection head tailored for small targets. Furthermore, the integration of positional information and surface features enables the method to achieve real-time monitoring and non-destructive early warning of fish surface abnormalities. The proposed method enhances precision in small target detection and achieves high accuracy in discerning subtle differences among detection targets. In real aquaculture settings, it can reach an average speed of 106 FPS with a detection accuracy of 98%. Although this study has been designed to meet the specific needs of zebrafish scientific research, it is highly generalisable and can be applied to the real-time detection of underwater surface abnormalities in a range of fish species in aquaculture.

## 1. Introduction

The zebrafish (*Danio rerio*) has emerged as a prominent model organism across a range of scientific fields, including medicine, developmental biology, genetics, toxicology and neurobiology. This prominence is largely attributed to its small size, rapid development, transparent embryos and a genome with a high similarity to humans (∼70%) [1]. Zebrafish are susceptible to a variety of diseases, among which infectious diseases pose the most severe threat, frequently leading to large-scale epidemic outbreaks [2]. Infected zebrafish commonly exhibit symptoms such as ulcerative bleeding on the body surface, congestion around the head, abnormal scales, congested gill filaments, tail corrosion, red or white discoloration, emaciation, spinal curvatures, and behavioural changes like slow swimming, solitary behavior, and resting at the bottom. As the application of zebrafish in scientific research expands, the need for non-invasive health monitoring has become increasingly evident. The condition of the body surface of zebrafish is a principal indicator of their health status. Early detection of surface abnormalities caused by diseases or other physiological stressors is essential for maintaining health and welfare of zebrafish, as well as for ensuring the validity of experimental results. Traditional techniques for diagnosing fish diseases encompass water quality analysis, behavioral observation, examination of the fish’s body surface, and microscopic examination [3, 4]. However, these conventional detection techniques rely on manual observation or the expertise of trained professionals, which can be time-consuming and labor-intensive nature of the process, prone to error, subject to personal subjective bias, and challenging to implement for continuous 24-hour surveillance [5]. These limitations may lead to missed sudden abnormalities, incomplete observational records, and hinder subsequent data statistics and analysis.

In recent years, methods based on machine learning and computer vision have begun to be applied to fish health management [6], where they are used to predict and even identify fish diseases by monitoring water quality, fish behavior or body surface abnormalities. These approaches offer significant advantages, including low cost, high efficiency, and non-invasiveness, while enabling continuous, automated monitoring without temporal constraints, thus presenting a viable alternative to traditional manual inspection [7]. For example, modern zebrafish recirculating aquaculture systems commonly use automated technologies to monitor and control environmental conditions, as well as to detect and regulate water quality through sensors and other devices, thereby preventing fish diseases [8, 9]. Additionally, Nayan et al. [10] employed gradient boosting models to analyze water quality, resulting in high accuracy in detecting fish diseases related to water quality. Fish movement behavior is also an important indicator of fish health status [4]. Pinkiewicz et al. [11] developed a computer vision system that can continuously monitor the movement trajectory of Atlantic salmon in real time to detect abnormal behaviors. Zhao et al. [12] proposed an innovative approach that integrates the modified motion influence map and recurrent neural network (RNN) to detect, locate, and identify localized abnormal behaviors within intensive aquaculture environments. Li et al. [13] achieved accurate tracking of a single abnormal fish by improving YOLOv5 and combined it with image fusion techniques, while Wang et al. [14] achieved multi-target tracking of abnormal fish by optimizing YOLOv5s model and combining it with a single-object tracking algorithm, SiamRPN++. Body surface detection is the most direct means of diagnosing fish diseases and can monitor fish health more accurately than monitoring water quality and fish behavior. Automated monitoring fish body surface using computer vision technologies has the potential to significantly reduce the time required for the diagnosis of contagious and hazardous fish diseases [15]. As some attempts, a fish disease detection method based on K-means and C-means fuzzy logic clustering has been used for automated detection and classification of body surface diseases in freshwater fish [16]; machine learning models (SVM) have also been used to determine and classify body surface abnormalities of salmon [17], and classification algorithms such as Random Forests can detect affected disease-affected areas through image segmentation to more accurately identify fish diseases [18].

Advancements in deep learning have led to substantial improvements in the accuracy of image analysis and recognition. Algorithms based on deep convolutional neural networks (DCNN) have proven effective in detecting ulcers and white spots on the surface of fish [19]. Furthermore, models trained through transfer learning and convolutional neural networks (CNN) have exhibited the capability to classify three distinct types of abnormal body surfaces in grouper fish [20]. The modification of networks and the introduction of attention mechanisms further enhance the computer’s ability to process subtle differences and complex backgrounds. A CNN incorporating dropout layers and contrast-limited adaptive histogram equalization (CLAHE) has been used to more accurately identify lice and wounds on farmed fish [21]. The addition of supplementary detection layers to the YOLOv4 PANet has been demonstrated to facilitate the effective detection of tiny surface parasites in images captured by stereo microscopes [22]. The enhanced YOLOv5-s model, integrated with the CBAM (Convolutional Block Attention Module) attention mechanism, not only enhances the detection of smaller targets but also addresses the challenges of deep feature extraction in complex settings [23]. A CNN constructed by combining the Normalization-based Attention Module (NAM) with YOLOv7 was able to detect fish with symptoms of spring viremia of carp (SVC) [24]. In addition, the model constructed by adding ByteTrack, Long Short-Term Memory (LSTM), and fuzzy inference to YOLOv8 was able to synthesize multimodal information from body surface, water quality, and fish behavior to provide early warning of nocardiosis in largemouth bass [25].

However, it is notable that the aforementioned studies, along with most publicly accessible databases, such as the SalmonScan dataset [26], are predominantly based on images of fish that have been removed from their natural aquatic environment or placed in specialized apparatus, which limits the scope and potential applications of such studies. Moreover, most algorithms focus on only a single aspect of fish abnormalities. Due to the complexity of fish living environment and the diversity of fish species, the identification and diagnosis of fish body surface abnormalities are more difficult compared to water quality monitoring and fish behavioral studies, and have not yet been widely applied. For zebrafish, which serve as critical experimental models, the ability to conduct continuous, non-disruptive underwater monitoring is essential. Water quality monitoring is limited in the type of fish diseases it can warn of, and tracking zebrafish movements in dense schools faces problems such as individuals masking each other [27]. Fortunately, zebrafish are typically reared in 5-10 L transparent tanks, where clear underwater images are relatively easy to obtain. Therefore, multimodal information fusion, which is mainly based on body surface information and complemented by information such as fish position, is important for early warning and effective management of zebrafish diseases. However, the detection of zebrafish body surface still faces many challenges. First, the high stocking density (approximately 5 fishes per liter) and rapid swimming behavior of zebrafish can lead to occlusion and overlap during imaging, resulting in blurry images or partial loss of information. Second, the small size of zebrafish makes the subtle differences in surface abnormalities more susceptible to interference in a noisy rearing environment [28]. Accurate identification and localization of these details require high-resolution images and sophisticated detection models. Moreover, images taken from the edges or bottom of the tank may suffer from distortion, deformation, and blurred edges due to the camera’s distance and angle, reducing the effective features for detection. The requirement for real-time detection also necessitates the processing of a large volume of image data. Consequently, the detection of surface abnormalities in zebrafish necessitates a higher degree of accuracy, sensitivity, and image processing speed from the detection model.

The current mainstream real-time object detectors are the YOLO family series [29], which are widely used in scenarios requiring high frame rates, such as real-time video analysis. From YOLOv1 to YOLOv8, the YOLO models have continuously improved in terms of speed and accuracy [30]. YOLOv8, as a SOTA (State-of-the-Art) model, has made significant advancements in the field of object detection, offering higher performance, greater flexibility, and ease of extension and deployment [31]. This study proposes a method based on the YOLOv8 framework (ESC-YOLOv8-seg). The approach involves enhancing image denoising preprocessing, refining and optimizing the YOLOv8 model, and strengthening the detection of small targets. It aims to obtain more comprehensive spatiotemporal information about small targets and integrate the positional information and body surface features of fish for real-time monitoring and early warning of surface abnormalities. For this study, zebrafish infected with *Aeromonas hydrophila* (Ah) were used as the experimental subjects to acquire image data. Ah is a major fish pathogen, and infected fish typically exhibit congestion in various parts of the body, including both sides, fin bases, back, abdomen, tail. In severe cases, the lesions can lead to decay [32]. In zebrafish, a range of symptoms have been observed, such as skin ulcers, damage, rupture, local redness, and peeling, indicating a diverse range of surface abnormalities. To validate the effectiveness of this method, an online detection platform (Fishsitter, http://bioinfo.ihb.ac.cn/fishsitter/) has been established. The platform has confirmed the effectiveness and good performance of the method. This method also provides a reference for the development of real-time underwater detection technology for other fish species.

## 2. Materials and methods

### 2.1 Fish and video acquisition

Prior to initiating the experiment, Ah was inoculated into Tryptone Soy Broth (TSB) medium and incubated at 32°C with shaking at 150 rpm for 24 hours. The bacterial concentration was determined using a UV-Vis spectrophotometer. Zebrafish were divided into control and treatment groups and reared in tanks at a density of approximately seven fish per liter. The experiment lasted for two weeks, during which the fish were fed according to standard procedures. The control group was maintained in aerated water, while the treatment group was exposed to aerated water supplemented with a bacterial suspension, The water and bacterial suspension were replaced daily to maintain the Ah concentration at a predetermined sub-lethal level of 2×10^5 CFU/mL, as established in preliminary experiments. The treatment group consisted of 250 zebrafish, aged 2-4 months. Imaging began three days post-infection, when the diseased fish in the treatment group started to exhibit surface abnormalities, social withdrawal, and lethargy. Both affected individuals in the treatment group and healthy individuals in the control group were imaged.

Imagery was collected at varying resolutions for two distinct scenarios. In the first scenario, single-fish imaging was conducted using a PixelLink camera to capture close-up images of an individual fish confined in a small container at a rate of 80 frames per second (FPS) (Figure 1A). These images were of high resolution (2448 × 2048 pixel), and provided clear visual details. For the single-fish scenario, 200 frames were captured per fish prior to the onset of disease, and 200 frames were captured daily, split between morning and evening sessions, throughout the disease period until the fish’s death. This process typically spanned two days from the first appearance of surface abnormalities to the fish’s demise. In the second scenario, schooling-fish imaging was performed to reflect the actual rearing conditions of zebrafish. One to three fishes exhibiting signs of disease were housed in a standard PVC aquarium alongside a larger number of healthy control fish. The imaging equipment comprised a Xiaomi camera and a Huawei smartphone, positioned laterally to the aquarium. Over the course of 14 days, imagery was collected multiple times at a rate of 30 FPS with a resolution of 1920 × 1080 pixels. In these images, the fish appeared smaller and the background noise was more complex, reflecting the challenges of imaging in a natural, dynamic environment.

**Figure 1:**
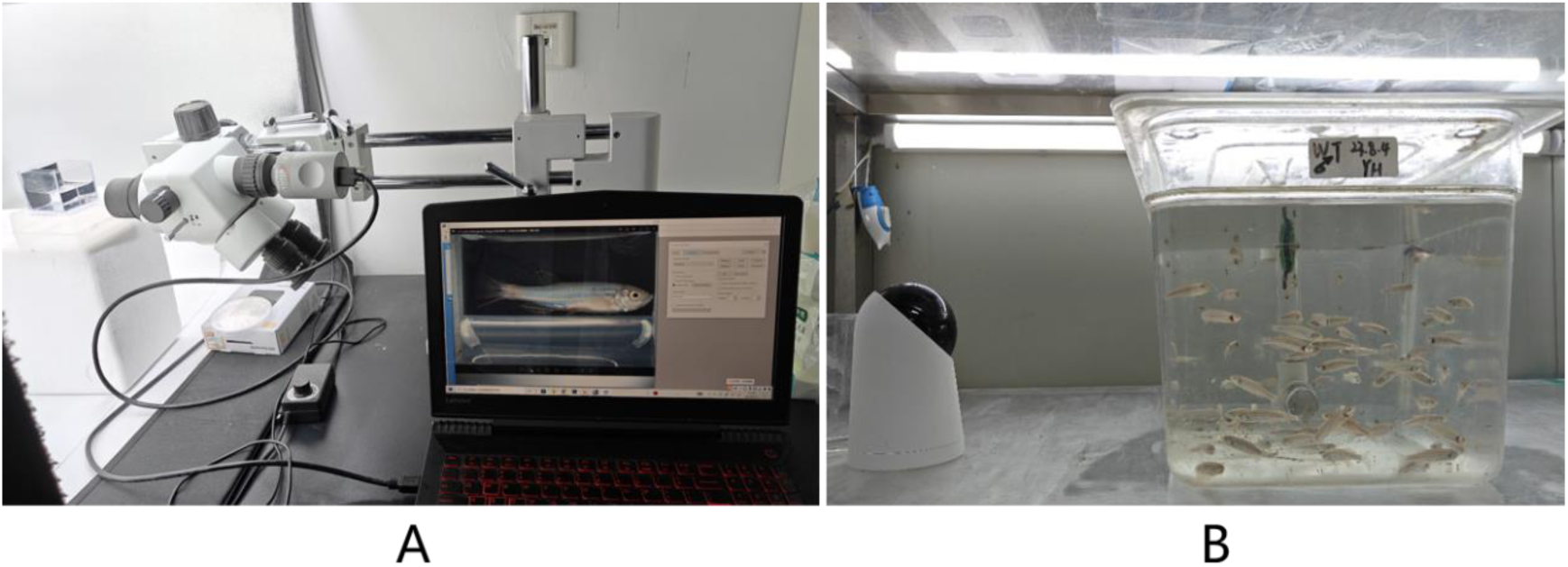
Image acquisition scenarios. A. Scene depicting the capture of high-quality images of a single zebrafish. B. Scene depicting the capture of a group of fish under normal farming conditions.

### 2.2 Data acquisition

#### 2.2.1 Image acquisition

We manually selected frames from the video sequences that exhibited various surface abnormalities, including scale loss, base of tail congestion, back congestion, lateral body congestion, abdominal congestion, and surface ulcers (Figure 2). This selection was made to enhance the diversity and representativeness of the experimental dataset. A total of 10,868 were selected from the single-fish videos, and 2,457 images from schooling-fish videos. Subsequently, the fish bodies in these images were manually annotated with instance segmentation using LabelMe [33], and the annotations distinguished between healthy and diseased fish.

**Figure 2:**
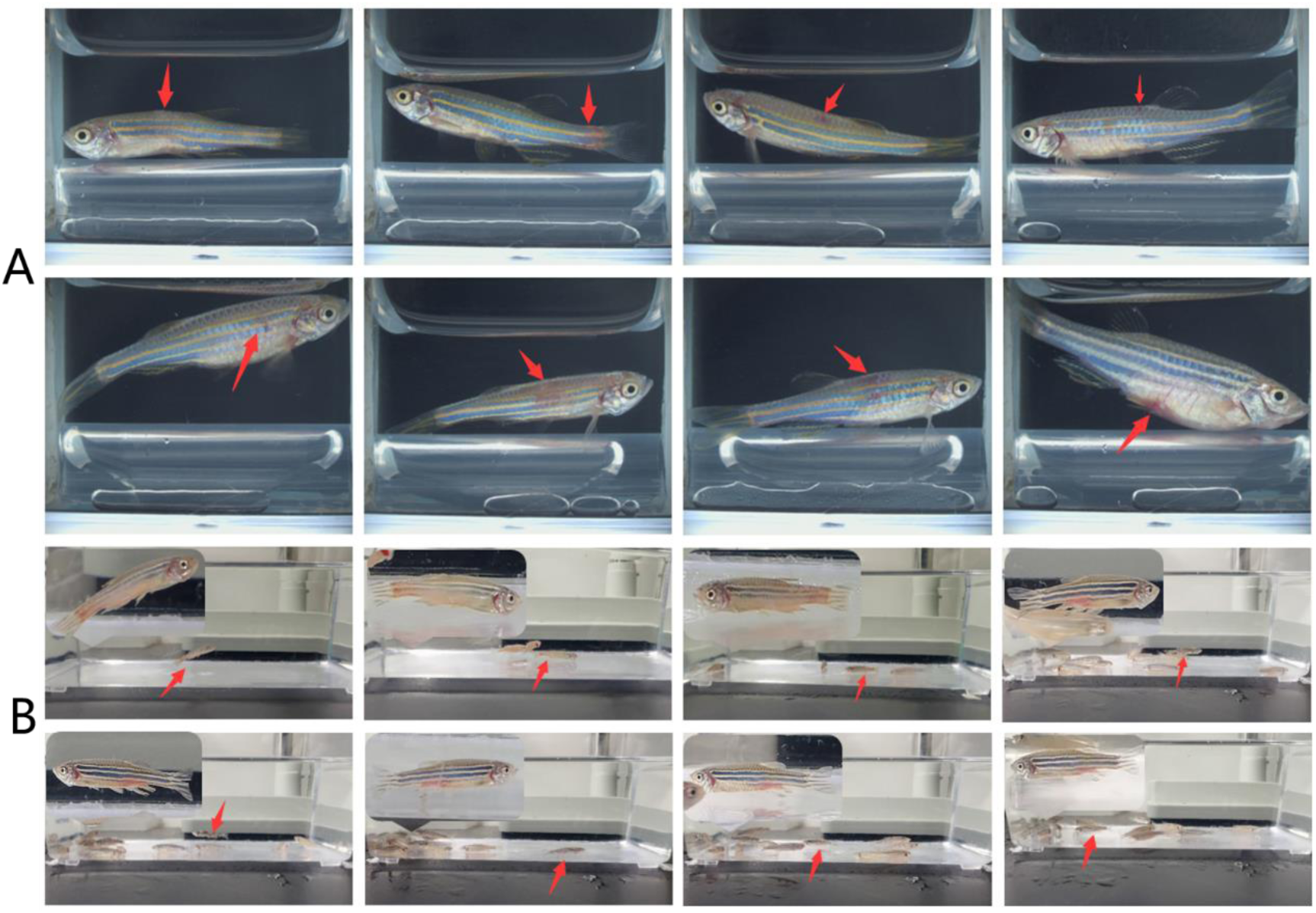
Schematic of surface abnormalities in zebrafish. A. Images of a single fish, with abnormal areas indicated by red arrows. B. Images of schooling fish, with diseased fish indicated by red arrows and an enlarged view of the diseased fish in the top-left corner.

#### 2.2.2 Image pre-processing

In response to the low contrast and complex noise characteristics of the schooling-fish images, we employed methods of contrast enhancement and bilateral filtering [34] for image preprocessing. Contrast enhancement improves the visibility of details and textures in images, thereby enhancing the features of small targets and improving recognition accuracy. However, this process may also introduce additional noise (Figure 3A-B). Therefore, subsequent denoising is necessary to reduce noise interference and further enhance the visibility of small targets (Figure 3C).

**Figure 3:**
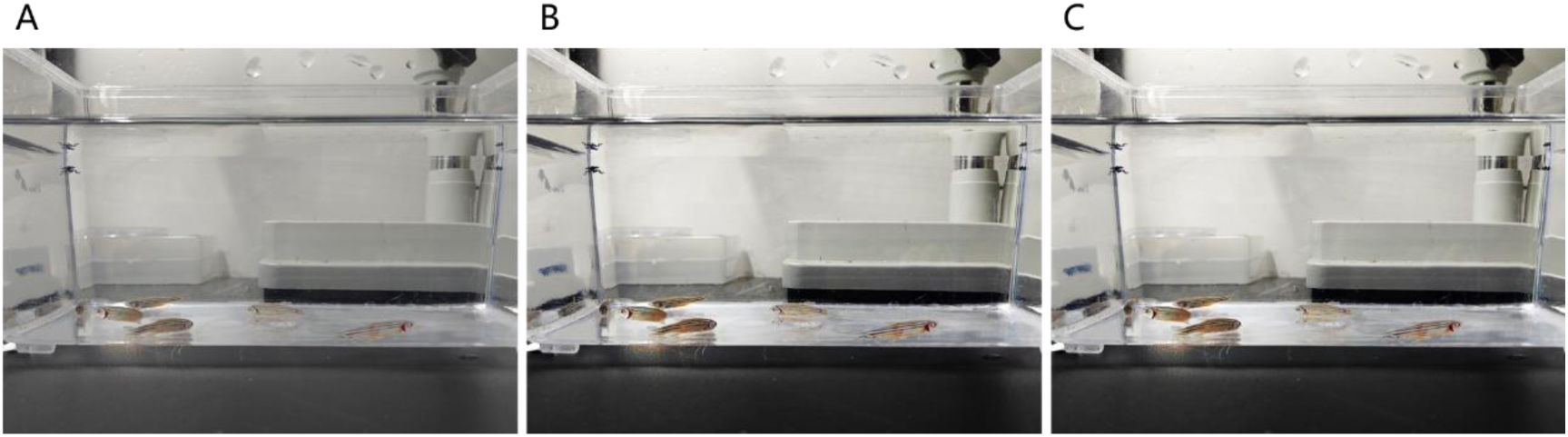
The impact of pre-processing on an image of school fish under normal farming conditions. A. the original image. B. The image after contrast enhancement processing. C. The image after denoising with bilateral filtering.

We constructed three datasets from all the collected images. The first dataset consists of the original schooling-fish images, some of which are of suboptimal quality. After applying image preprocessing techniques, the first dataset becomes the second dataset. The second dataset is then combined with single-fish images to form the third dataset. Each dataset is divided into two sections: one for training the model and the other for validating and testing the model (Table 1).

**Table 1:** Classification of the collected images.

| Data sets | Number of training<br>images | Number of validation<br>and test images | Total<br>images |
| --- | --- | --- | --- |
| Raw schooling-fish images | 1960 | 497 | 2457 |
| Preprocessed schooling-fish images | 1960 | 497 | 2457 |
| Mixed images | 10660 | 2665 | 13325 |
| single-fish images | 8700 | 2168 | 10868 |
| preprocessed schooling-fish images | 1960 | 497 | 2457 |

During the model training phase, we applied various data augmentation techniques to the training set images. These techniques included random cropping, rotation, and scaling, which serve to enrich the training data. This approach addresses the substantial need for labelled data when training deep learning models, and enhance the algorithm’s generalization capabilities and robustness.

### 2.3 Proposed method to identify abnormal surface of zebrafish

The ESC-YOLOv8-seg proposed in this study is based on the YOLOv8-seg network framework, which is implemented based on YOLACT [35]. The model architecture is illustrated in Figure 4. The input image (Figure 4A) is initially processed through the backbone network (Figure 4B) to extract features, which are subsequently merged with features of varying sizes by the FPN (Feature Pyramid Network) (Figure 4C) before entering two parallel branches. The detection branch (Figure 4D) outputs the class, bounding box coordinates (x, y, w, h), and k mask coefficients (either -1 or 1) for each target object. The segmentation branch (Figure 4E) generates k prototypes for the current input image. For each target object, the k mask coefficients are multiplied with the k prototypes, and the sum of all results (Figure 4F) yields the instance segmentation result, which is then outputted (Figure 4G).

**Figure 4:**
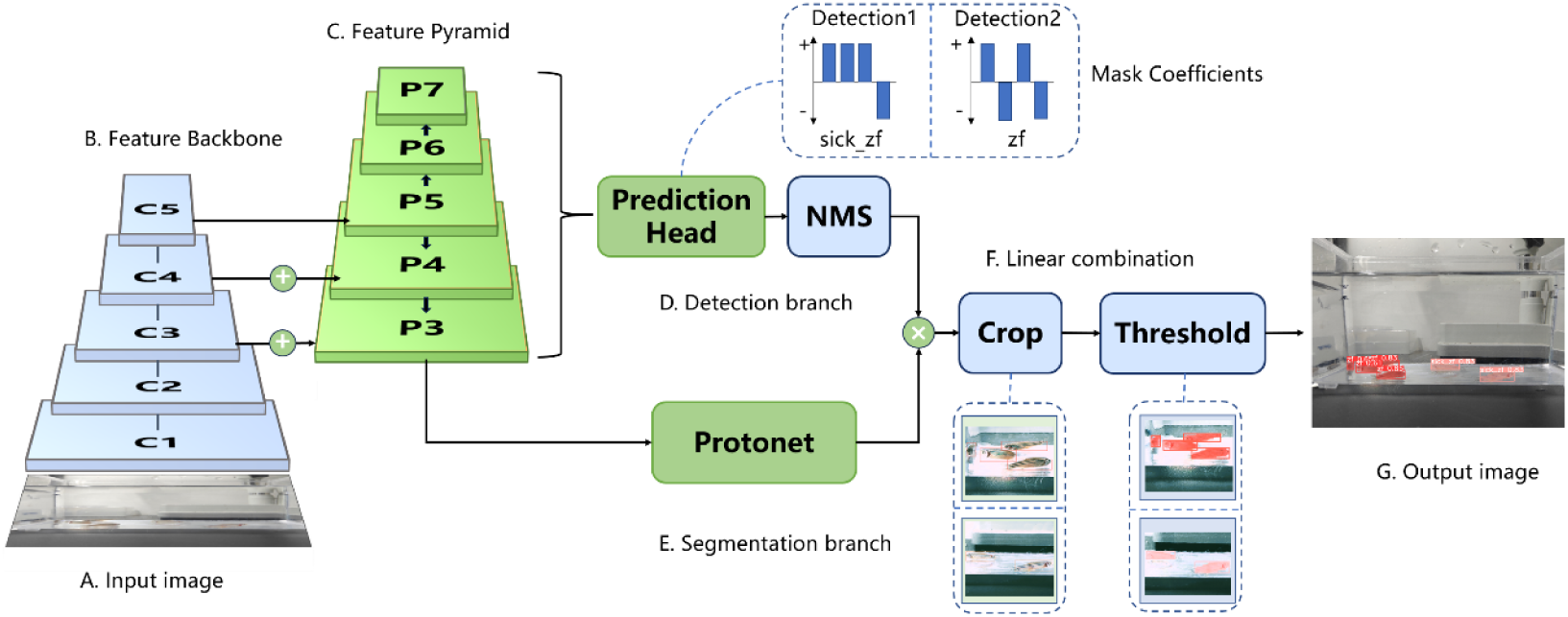
A detailed diagram for the network structure of YOLACT. A. Input image. B. Backbone. C. Feature Pyramid. D. Detection branch. E. Segmentation branch. F. Linear combination. G. Output image.

The ESC-YOLOv8-seg model introduces several key improvements to the YOLOv8-seg framework (Figure 5A-C). Firstly, the Efficient Multi-scale Attention (EMA) module [36] was incorporated between the C2f_Faster module and the Detect head (Figure 5B, Supplementary Figure 1). Subsequently, in order to address the issue of small targets being easily missed, the original SPPF module at the lowest layer of the backbone was replaced with the SPPELAN module [37] (Figure 5A, D). This module integrates the Spatial Pyramid Pooling (SPP) with a local attention mechanism to capture finer target features at different scales. Additionally, the bottleneck in the C2f module was replaced with a FasterBlock, and the modified C2f module was renamed the C2f_Faster module [38] (Figure 5A-B, F-G). Furthermore, a dedicated detection head, designated as P2, was developed within the Detect layer to enhance the identification of small and tiny-sized objects (Figure 5C, E).

**Figure 5:**
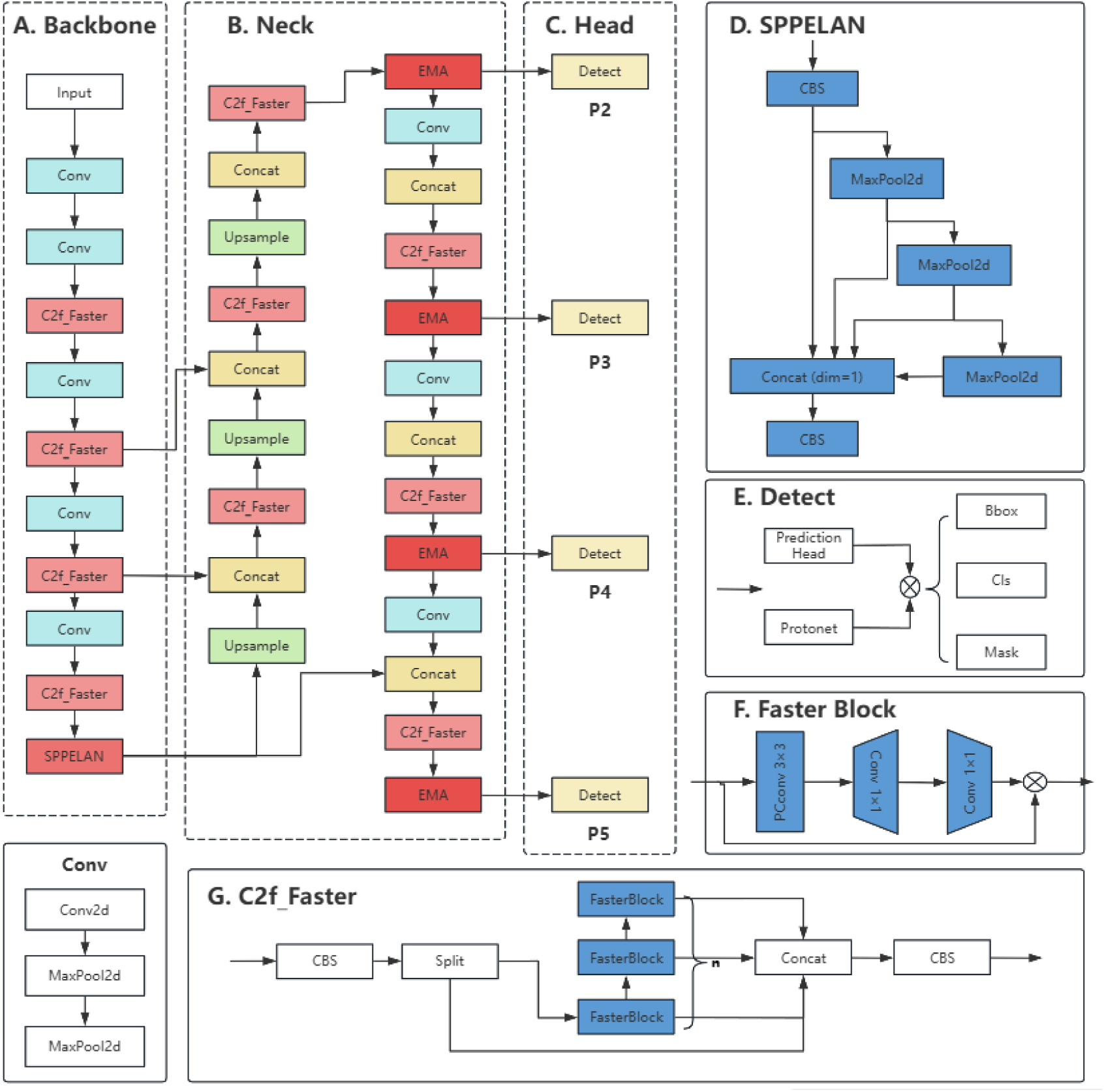
The schematic representation of the network architecture of ESC-YOLOv8-seg and its constituent subcomponents. A. Backbone network. B. Feature fusion Neck. C. Detection Head. D. SPPELAN. E. Detect. F. Faster Block. G. C2f_Faster.

In the initial stages of the detection process, the video or image data is subjected to a series of preprocessing steps prior to being fed into the YOLOv8 network. Subsequently, the input images are subjected to a process of feature extraction within the backbone (Figure 5A). Following this, feature fusion and enhancement are performed in the Neck (Figure 5B), where the extracted features are integrated and refined. Finally, within the Head, four Detect modules (Figure 5C) are employed to detect zebrafish targets of varying sizes, make decisions, and produce the corresponding outputs.

#### 2.3.1 Efficient Multi-Scale Attention Module

In the early stages of disease, zebrafish display minimal alterations in surface appearance when compared to healthy individuals. Nevertheless, some diseased fish may display behavioural changes, such as a reduction in swimming activity or a tendency to rest at the bottom of the tank. As the disease progresses and surface abnormalities become apparent, affected fish often isolate themselves from the group, highlighting the importance of positional information for target detection. Coordinate Attention (CA) represents an attention mechanism that incorporates positional data into channel attention, thereby effectively integrating spatial coordinate information into the generated attention maps. This integration enhances the model’s capacity to accurately locate and identify objects of interest [39]. However, the limited receptive field of the 1×1 convolution employed in CA’s network configuration hinders the modelling of local cross-channel interactions and the utilisation of contextual information, thus neglecting the significance of interactions between disparate spatial locations.

EMA is a non-reductive, efficient multi-scale focusing method that improves upon the sequential processing approach of CA by placing two distinct convolutional kernels within parallel sub-networks [36]. One sub-network employs a 1 × 1 convolutional kernel, operating similarly to CA, while the other uses a 3 ×3 convolutional kernel (Supplementary Figure 1). The parallel sub-structure allows the network to circumvent extensive sequential and deep processing. As a result, EMA not only retains more precise spatial location information for regions of interest in the final output, but also facilitates the network in extracting location information from feature maps, which is crucial for the accurate detection of targets based on the position of the fish bodies.

#### 2.3.2 SPPELAN

SPP is a pooling strategy introduced in YOLOv3 that captures a more comprehensive set of fine-grained features through multi-scale pooling operations and feature map processing. This enhances the receptive field of the model, thereby enabling the capture of both local and global information about objects, and enhances the model’s robustness when dealing with targets of varying scales and deformations [40]. These capabilities are particularly important for recognizing small targets like zebrafish, which occupy only a small area on the feature maps. The incorporation of fine-grained features and local information improves detection accuracy. Nevertheless, the computational cost of SPP is considerable, which can slow down the model’s inference speed.

The Efficient Layer Aggregation Network (ELAN) employs a layer aggregation strategy to integrate features from various layers, allowing the model to capture a wider range of contextual and detailed information, thus enriching and diversifying the feature representation [41]. Inter-layer feature fusion strengthens the model’s expressiveness and robustness, making it more reliable for detecting targets, especially small ones, in complex backgrounds, under occlusions, noise, and other interferences. Furthermore, ELAN’s lightweight design also reduces computational complexity and accelerates inference speed when integrated into a model.

SPPELAN, as illustrated in Figure 5D, is a combination of SPP and ELAN, designed to enhance the efficacy of object detection by leveraging the strengths of both components [37]. In the YOLOv8 architecture, the SPPF module has been replaced with the SPPELAN module at the base of the backbone (Figure 5A). By merging multi-scale feature extraction with efficient layer aggregation, SPPELAN not only improves the detection accuracy of small targets like zebrafish but also significantly enhances detection speed, making it suitable for a variety of real-time detection applications. For fast-moving small targets, SPPELAN provides precise and efficient detection and tracking.

#### 2.3.3 FasterNet Bolck

Zebrafish are highly sociable creatures that typically live and move in groups, leading to complex backgrounds and substantial occlusions during video detection procedures. This can cause convolutional neural networks (CNNs) to extract redundant features and noise, hindering the model’s recognition accuracy. Convolution operations require filtering at every location, and stacking of multiple layers in CNNs add to the model’s complexity, thereby demanding more computational resources. Eliminating superfluous features and reducing model complexity can enhance the accuracy and speed of target detection. FasterNet introduces a novel partial convolution (PConv) that systematically applies regular convolutions on selected input channels without affecting others, taking advantage of the redundancy in feature maps [7]. Moreover, it incorporates a point-wise convolution (PWConv) based on PConv to efficiently utilize information across all channels. Each FasterNet block comprises a PConv layer, followed by two PWConv (or 1×1 Conv) layers (see Figure 5F). Together, they form an inverse residual block structure, wherein the middle layer has an expanded number of channels and also incorporates shortcut connections to reuse input features. In this study, an improvement was made to the C2f module in the YOLOv8-seg network by replacing the bottleneck with a FasterNet block, creating a new structure called C2f-faster. This modification is aimed at feature extraction and processing for zebrafish target detection, maintaining a certain receptive field range and nonlinear representation capability while reducing the parameters and computational complexity, as illustrated in Figure 5G.

### 2.4 Evaluation of abnormal body identification results

Accurate target identification and classification play a crucial role in the real-time detection of abnormal fish. By default, YOLOv8 employs the Complete Intersection over Union (CIoU) as the loss function for bounding box regression. A prediction is deemed correct when the Intersection over Union (IoU) value exceeds a certain threshold, typically 0.5. The IoU values are computed as delineated in Equation (1):

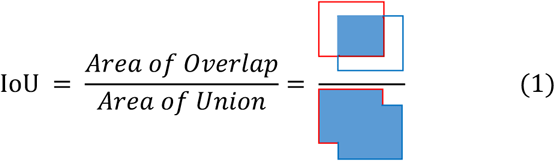

We employ a confusion matrix to assess the performance of our method. The confusion matrix evaluates the accuracy of the classifier based on four fundamental metrics: true positives (TP), true negatives (TN), false positives (FP), and false negatives (FN). Additional classification metrics, such as Accuracy, Precision, and Recall, can be derived from the confusion matrix. These metrics are calculated as follows in Equation (2):

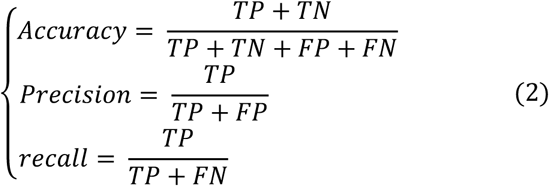

Accuracy is the proportion of zebrafish with correctly judged body surface conditions out of the total number of observed zebrafish. Precision indicates the ratio of correctly identified zebrafish body surface conditions to the sum of correctly and incorrectly identified conditions. Recall represents the ratio of correctly identified zebrafish body surface conditions to the total number of conditions that were either correctly or not identified.

The Average Precision (AP) is calculated as delineated in Equation (3).

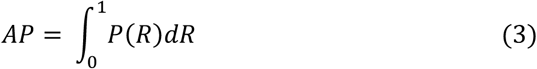

The mean Average Precision (mAP) at an Intersection over Union (IoU) threshold of 0.5, denoted as mAP@0.5, is determined by computing the AP for the classification of normal and abnormal zebrafish body surfaces at an IoU of 0.5 and then averaging these values. The mAP@0.5:0.95 is calculated by setting the IoU threshold to range from 0.50 to 0.95 in increments of 0.05, computing the mAP for each IoU threshold, and subsequently averaging these values.

## 3. Results

### 3.1 Implementation details

The model architecture of this study is deployed on a DELL T3650 workstation with the following hardware specifications: an Intel Core i7-11700 processor (8 cores with a base frequency of 2.5 GHz and a turbo frequency of up to 4.9 GHz), 32 GB of RAM, an NVIDIA RTX A4000 graphics card, a 1 TB mechanical hard drive, and a 256 GB solid-state drive. The versions of PyTorch and CUDA in use are 1.12.0 and 11.3, respectively. An online testing platform has been developed, utilizing ESC-YOLOv8-seg as the backend framework and Django as the frontend framework.

### 3.2 Comparisons of popular models

Considering that ESC-YOLOv8-seg is based on YOLOv8-seg, we initially compared YOLOv8-seg with two other established models, namely Mask R-CNN [42] and YOLACT[35], using FPS and mAP as evaluation metrics. The models were trained and tested on raw schooling-fish images, with a batch size of 16 and 100 epochs. The results indicated that, in real zebrafish aquaculture settings, YOLOv8-seg not only had a significantly smaller model weight in comparison to YOLACT and Mask R-CNN, but also demonstrated the fastest detection speed, reaching 179 FPS, which is more than 35 times faster than Mask R-CNN (Table 2).

**Table 2:** Statistics of the results of the comparison of the three models.

| Models | mAP@0.5 |  | mAP@0.5:0.95 |  | mAP@0.5 |  | mAP@0.5:0.95 |  | FPS | weight |
| --- | --- | --- | --- | --- | --- | --- | --- | --- | --- | --- |
|  | (Box) |  | (Box) |  | (Mask) |  | (Mask) |  |  |  |
|  | healthy | sick | healthy | sick | healthy | sick | healthy | sick | -- | -- |
| Mask R-CNN | 0.855 | 0.933 | 0.702 | 0.592 | 0.875 | 0.895 | 0.590 | 0.630 | 5 | 244MB |
| YOLACT | 0.954 | 0.944 | 0.722 | 0.774 | 0.648 | 0.680 | 0.357 | 0.551 | 51 | 117MB |
| YOLOv8-seg | 0.968 | 0.994 | 0.791 | 0.915 | 0.968 | 0.954 | 0.661 | 0.742 | 179 | 6.45MB |

Note: healthy means normal zebrafish and sick means abnormal zebrafish

Regarding mAP values (Figure 6), YOLACT’s mask detection accuracy of was significantly lower than that both of Mask R-CNN and YOLOv8-seg, with YOLOv8-seg slightly outperforming Mask R-CNN. With regard to the segmentation of instances (masks), the mAP@0.5 value of YOLACT is 0.221 and 0.297 lower than that of Mask R-CNN and YOLOv8-seg, respectively. Furthermore, the mAP@0.5:0.95 value is 0.156 and 0.248 lower, respectively. Conversely, when detecting objects (boxes), the mAP value of Mask R-CNN is lower than that of YOLACT and YOLOv8-seg, with YOLOv8-seg also exhibiting superior performance to YOLACT.

**Figure 6:**
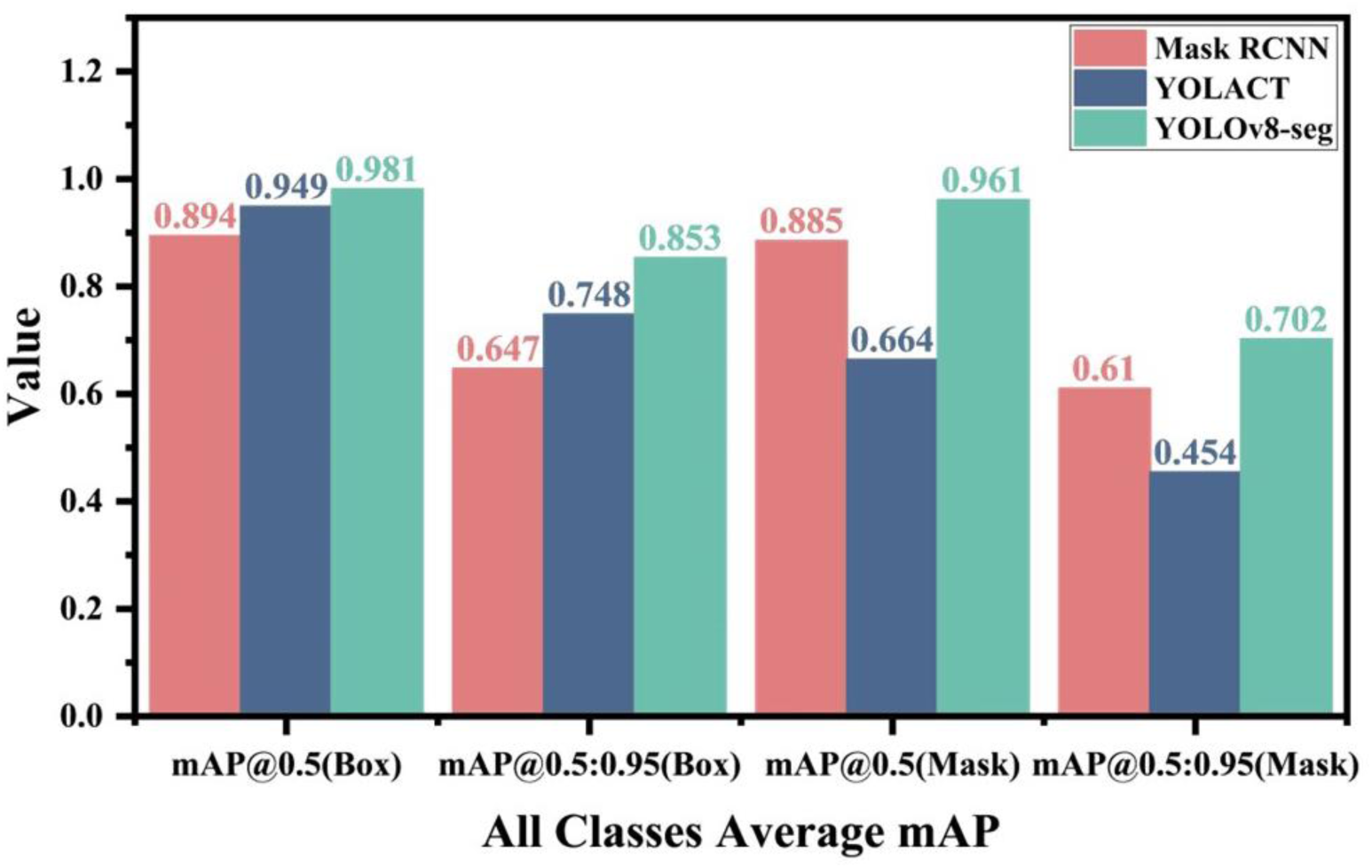
Mean Average Precision (mAP) values of three models following training on the Raw schooling-fish images dataset during detection testing.

The results presented above show that YOLOv8-seg outperforms Mask R-CNN and YOLACT in terms of both detection accuracy and speed, while also exhibiting the smallest model weight. In real recirculating aquaculture environments, it is crucial that a detection model demonstrates high accuracy, rapid processing speed, and a small weight for easy deployment in order to accurately identify abnormalities on fish body surfaces. Therefore, we decided to base our model for abnormal body surface detection model on YOLOv8-seg.

### 3.3 Detection performance of modified models

Building upon the YOLOv8-seg framework, we implemented refinements using CA, EMA, and SPPELAN, respectively. The modified models, along with the comprehensively enhanced ESC-YOLOv8-seg model, were trained and tested across a range of image datasets, with a batch size of 16 and a total number of training cycles (Epoch) set to 300. The performance of the models was then evaluated based on several metrics, including precision, recall, mAP, FPS, model size, detection speed, and the number of model parameters.

The modifications led to varying degrees of improvement in detection accuracy. However, there was an inverse relationship between model complexity and FPS, with FPS decreasing as model complexity increased (Table 3). When training and testing using raw schooling-fish images, the instance segmentation precision progressively improved with CA, EMA, and SPPELAN, with ESC-YOLOv8-seg exhibiting the best performance. The mAP@0.5 and mAP@0.5:0.95 values were found to be 0.021 and 0.105 higher, respectively, in comparison to those observed for YOLOv8-seg. Moreover, applying image contrast enhancement and denoising to these datasets resulted in an additional increase in mAP@0.5 and mAP@0.5:0.95 values by 0.023 and 0.104, respectively. Using preprocessed schooling-fish images generally elevated the detection accuracy of the models. Incorporating high-resolution single-fish images (mixed images) resulted in the best results for target instance segmentation, with mAP@0.5 and mAP@0.5:0.95 reaching 0.994 and 0.857, respectively. This represents an improvement of 0.033 and 0.112 over YOLOv8-seg, respectively.

**Table 3:** Recognition results of the models after various modifications across different datasets.

| Data | Models | mAP@0.5<br>(Mask) | mAP@0.5:0.95<br>(Mask) | FPS | layer |
| --- | --- | --- | --- | --- | --- |
| Raw schooling-fish images | YOLOv8-seg | 0.961 | 0.702 | 179 | 261 |
|  | YOLOv8seg + CA | 0.964 | 0.722 | 164 | 291 |
|  | YOLOv8-seg + EMA | 0.966 | 0.723 | 132 | 285 |
|  | YOLOv8-seg + SPPELAN | 0.972 | 0.737 | 146 | 263 |
|  | ESC-YOLOv8-seg | 0.982 | 0.807 | 106 | 377 |
| Preprocessed schooling-fish images | YOLOv8-seg | 0.965 | 0.710 | 182 | 261 |
|  | YOLOv8-seg + EMA | 0.972 | 0.734 | 136 | 285 |
|  | ESC-YOLOv8-seg | 0.988 | 0.814 | 110 | 377 |
| Mixed images | YOLOv8-seg | 0.961 | 0.841 | 180 | 261 |
|  | YOLOv8-seg + EMA | 0.974 | 0.843 | 144 | 285 |
|  | ESC-YOLOv8-seg | 0.994 | 0.857 | 112 | 377 |

Figure 7 illustrates the enhanced performance of several modified models. After training with three types of datasets, ESC-YOLOv8-seg achieved the highest precision and recall, both of which approached 1. This typically indicates that the model’s classification performance is excellent, with a strong ability to accurately distinguish between different classes of samples. With regard to the datasets, the preprocessed schooling-fish images outperformed raw schooling-fish images, while the mixed images demonstrated greater efficacy than the former. This highlights the critical importance of noise reduction preprocessing and the use of high-quality, single-fish data in improving model accuracy.

**Figure 7:**
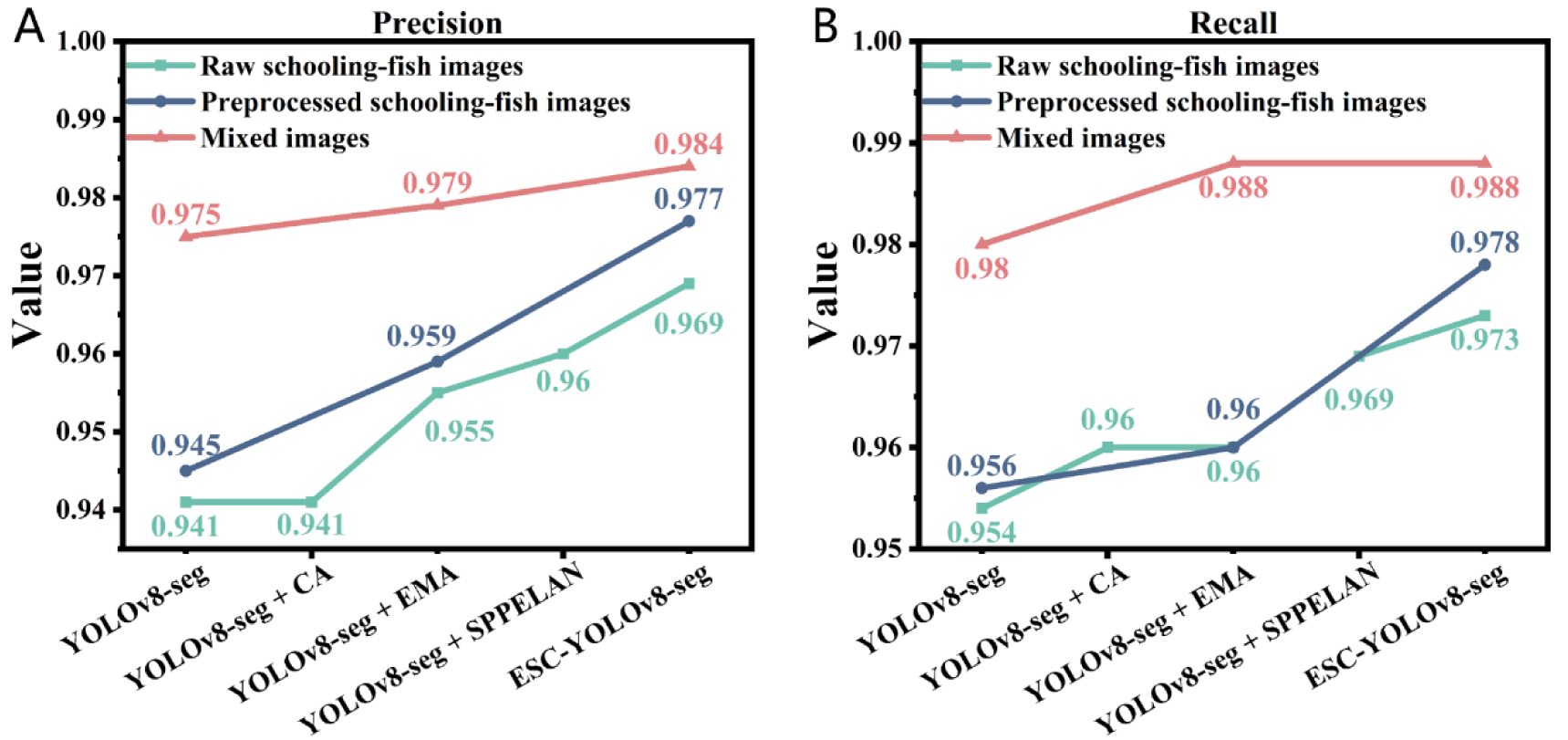
Results of various modified models upon completion of training on different datasets. A. Precision. B. Recall.

To mitigate the risk of overfitting, we validated the YOLOv8-seg and ESC-YOLOv8-seg models using test sets from two categories of data: raw schooling-fish images and mixed images. Compared to YOLOv8-seg, ESC-YOLOv8-seg reduced the number of FPs, where healthy zebrafish were incorrectly identified as sick, from 51 to 35 in the Raw schooling-fish images test set. The number of false negatives (FNs), where sick fish were incorrectly classified as healthy, decreased from 50 to 41. This improvement led to an increase in overall accuracy from 90.12% to 92.65% (Figure 8 A-B). For the mixed images test set, the number of FPs was reduced from 78 to 39, and the number of FNs from 58 to 20, resulting in an accuracy improvement from 95.75% to 98.25% (Figure 8 C-D). The confusion matrix demonstrates the significant effectiveness of the modifications that implemented in the model.

**Figure 8:**
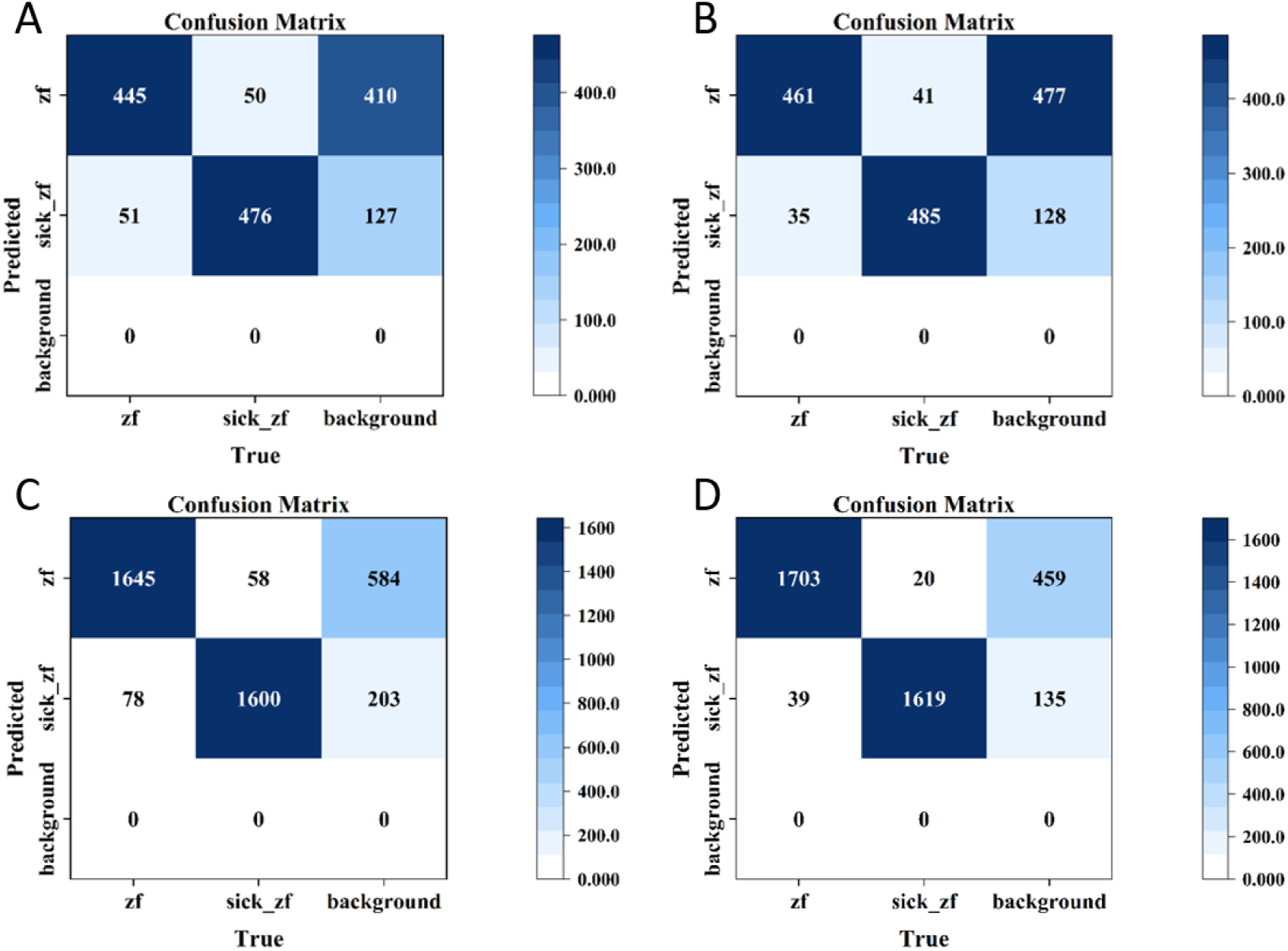
Confusion matrices. A-B. Detection results of YOLOv8-seg and ESC-YOLOv8-seg on the Raw schooling-fish images dataset. C-D. Detection results of YOLOv8-seg and ESC-YOLOv8-seg on the Mixed images dataset.

### 3.4 Model recognition results

The ESC-YOLOv8-seg model proposed in this study demonstrated a superior detection accuracy rate of 97.21%, exceeding the original YOLOv8-seg’s lower accuracy rate. Figure 9 illustrates the recognition outcomes of the images using the ESC-YOLOv8-seg model, with all sample images being correctly identified. Figure 10 provides a comparison of the recognition results obtained from the YOLOv8-seg and ESC-YOLOv8-seg models on the same set of images, with the incorrect identifications marked by yellow arrows. The YOLOv8-seg model exhibited a notable tendency towards erroneous recognition, with a total of four FPs and two FNs (Figure 10B). In contrast, the ESC-YOLOv8-seg model demonstrated a markedly superior accuracy, with only a single FP (Figure 10C).

**Figure 9:**
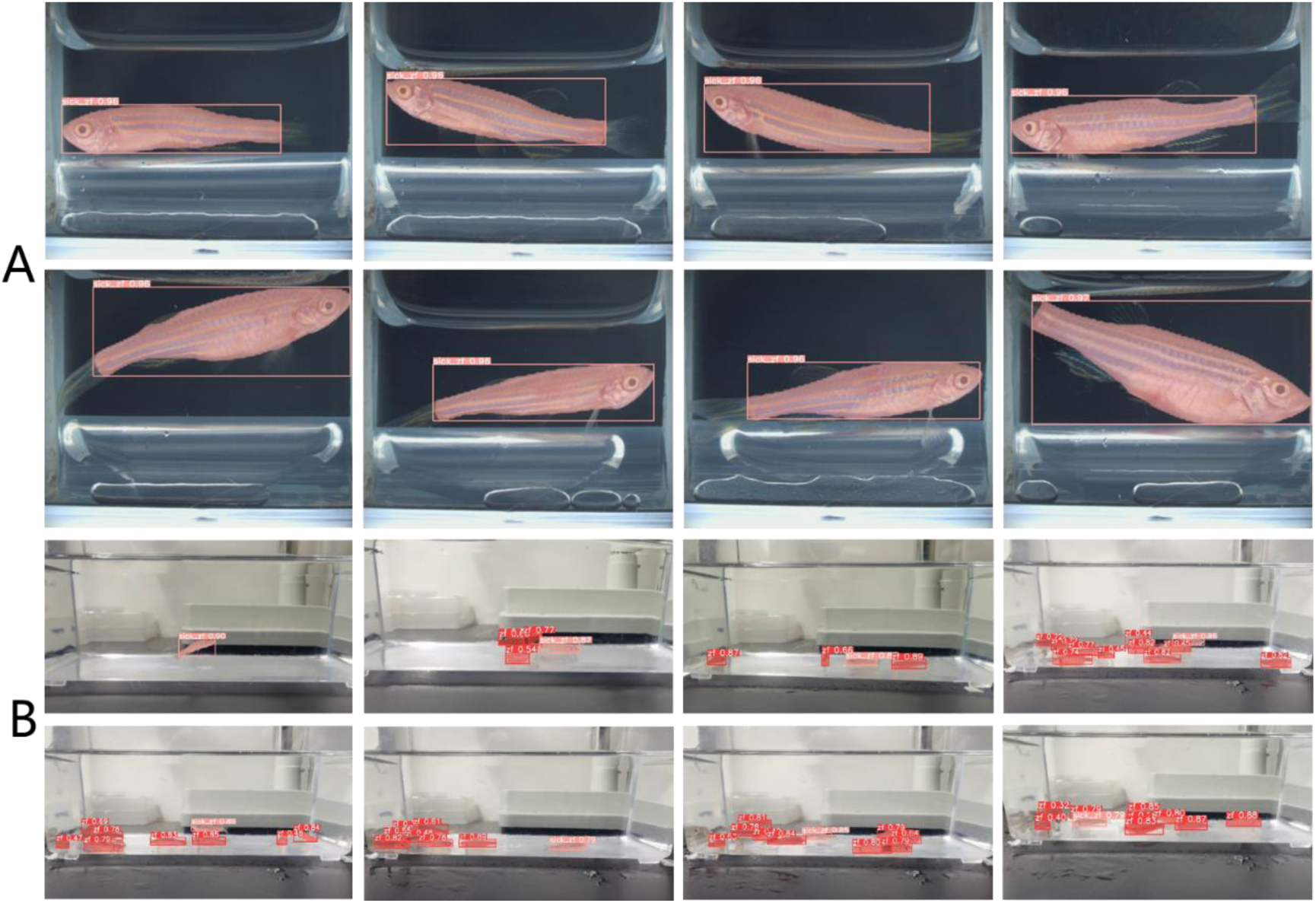
Recognition results of the sample images depicted in Figure 2. A. The detection results for the images of single fish. B. The detection results for the images of schooling fish.

**Figure 10:**
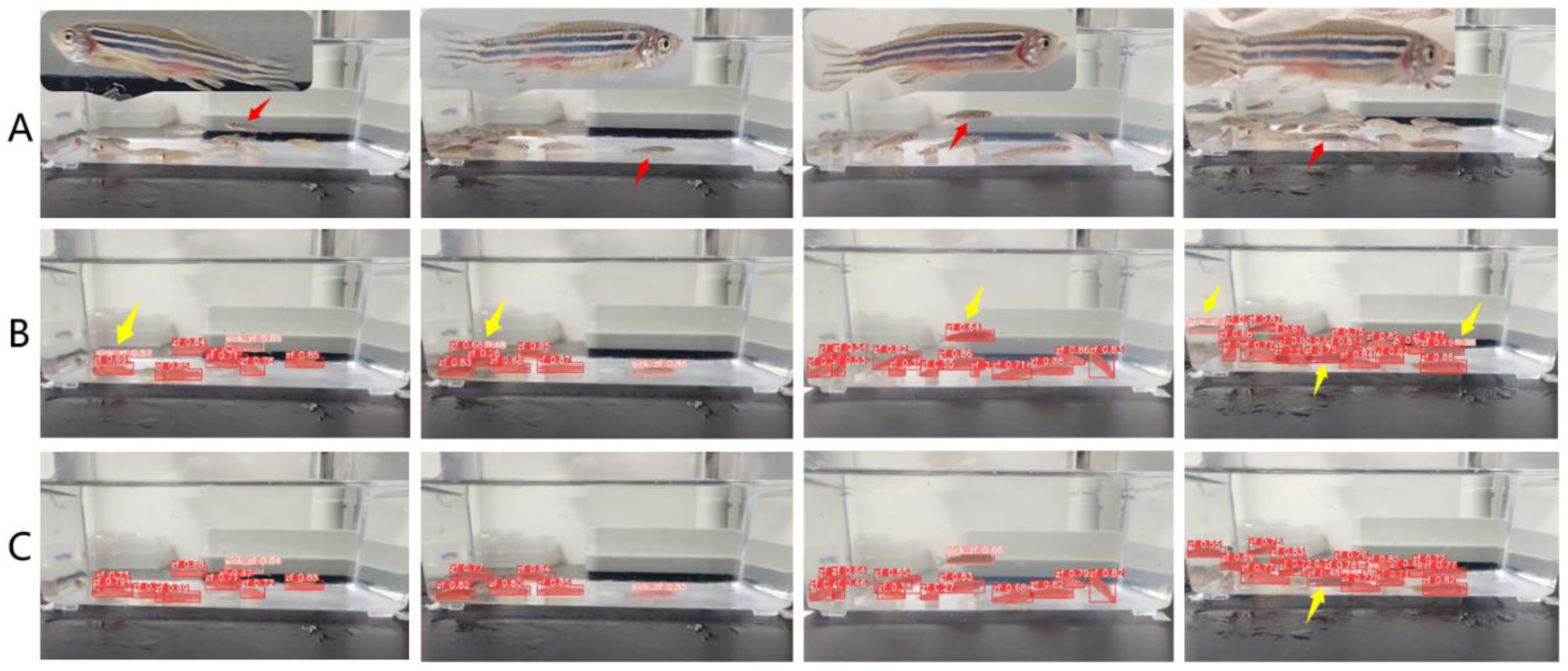
A comparison of the recognition results obtained from the YOLOv8-seg and ESC-YOLOv8-seg models. A. Original images of schooling fish, with diseased fish indicated by red arrows and an enlarged view of the diseased fish in the top-left corner. B-C. Recognition results of YOLOv8-seg and ESC-YOLOv8-seg, respectively. Yellow arrows indicate incorrect recognition results.

### 3.5 An attempt at surface localiztion of lesions

In light of the classification capabilities of the YOLOv8-cls framework integrated within

YOLOv8, we conducted a preliminary investigation into its potential application for lesion localization. For single-fish images depicting diseased zebrafish, we further processed the segmented fish body images, and performed a simple classification of the affected regions, categorizing them as caudal peduncle, abdomen, back, and pectoral fins. Out of the 10,868 single fish images, 4,777 depicted diseased zebrafish. Additionally, 1,000 randomly selected images of healthy zebrafish were included, resulting in a total of 5,777 images classified into five categories: ori_fish, sick_tail, sick_back, sick_stomach, and sick_fin, representing healthy zebrafish and fish with lesion in different body regions, respectively. Following the labelling of these images and subsequent training of the model, YOLOv8-cls proceeded to classify the segmented single fish images, providing the probability of each fish belonging to every classification category. The classification results could be integrated into the original detection data (Figure 11). To assess the model’s performance, a random sample of 567 images, not part of the training set, was selected for testing. The overall error rate was determined to be 8.56% (Table 4). In general, a higher predicted probability (*P*-value) is associated with a lower likelihood of classification error. In 72% (416/576) of the images, the maximum *P*-value of 1, with an error rate of only 0.96%, indicating a high level of reliability. However, when the maximum *P*-value was less than 1, the error rate increased significantly, suggesting that the lesions in such cases may be atypical and require cautious interpretation of the results.

**Figure 11:**
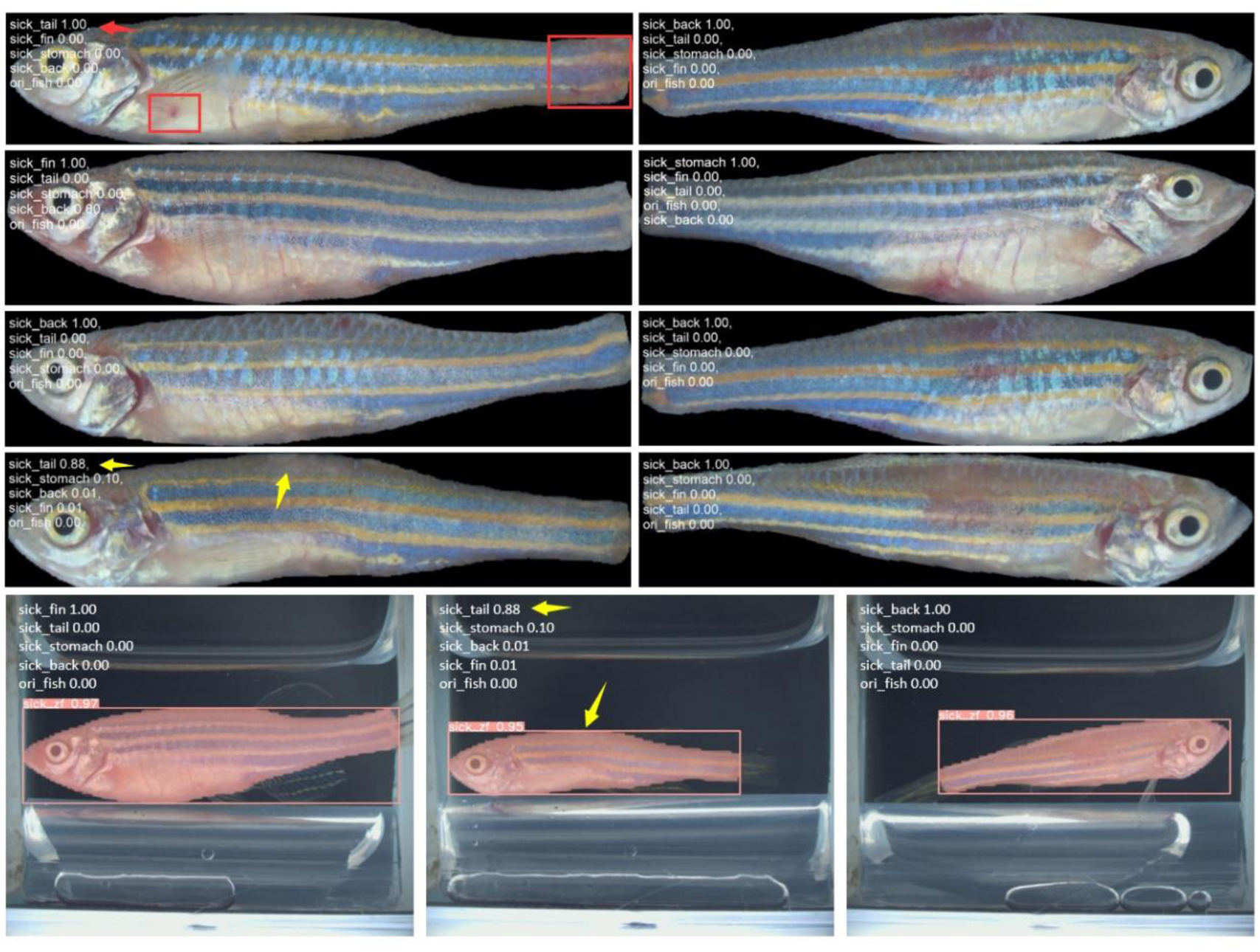
Classification results of segmented single diseased fish images and integrated recognition outcomes. Red boxes and arrows highlight the lesion locations and classification results, while the yellow arrows indicate the true lesion areas and erroneous recognition instances.

**Table 4.** Statistical summary of classification results for lesion locations in diseased zebrafish.

| Maximum classification probability | Total number | The expert checked the number of identification errors | Identification error rate |
| --- | --- | --- | --- |
| $P = 1$ | 416 | 4 | 0.96% |
| $1 > p \geq 0.9$ | 103 | 23 | 22.33% |
| $0.9 > P \geq 0.5$ | 44 | 19 | 43.18% |
| $P < 0.5$ | 4 | 2 | 50.00% |
| Total | 567 | 48 | 8.56% |

### 3.6 Functions of online platfrom

The ESC-YOLOv8-seg-based platform has the potential to facilitate health monitoring in zebrafish aquaculture systems. For real-time zebrafish monitor, it is essential to implement the methodological framework locally. To evaluate the efficacy of our approach, we have also developed a non-real-time online platform (Fishsitter, http://bioinfo.ihb.ac.cn/fishsitter/, Figure 12), which offers two interfaces for the detection testing of both images and videos. The help section of the platform provides users with access to the open-source code, installation instructions, and the experimental data used in this study.

**Figure 12:**
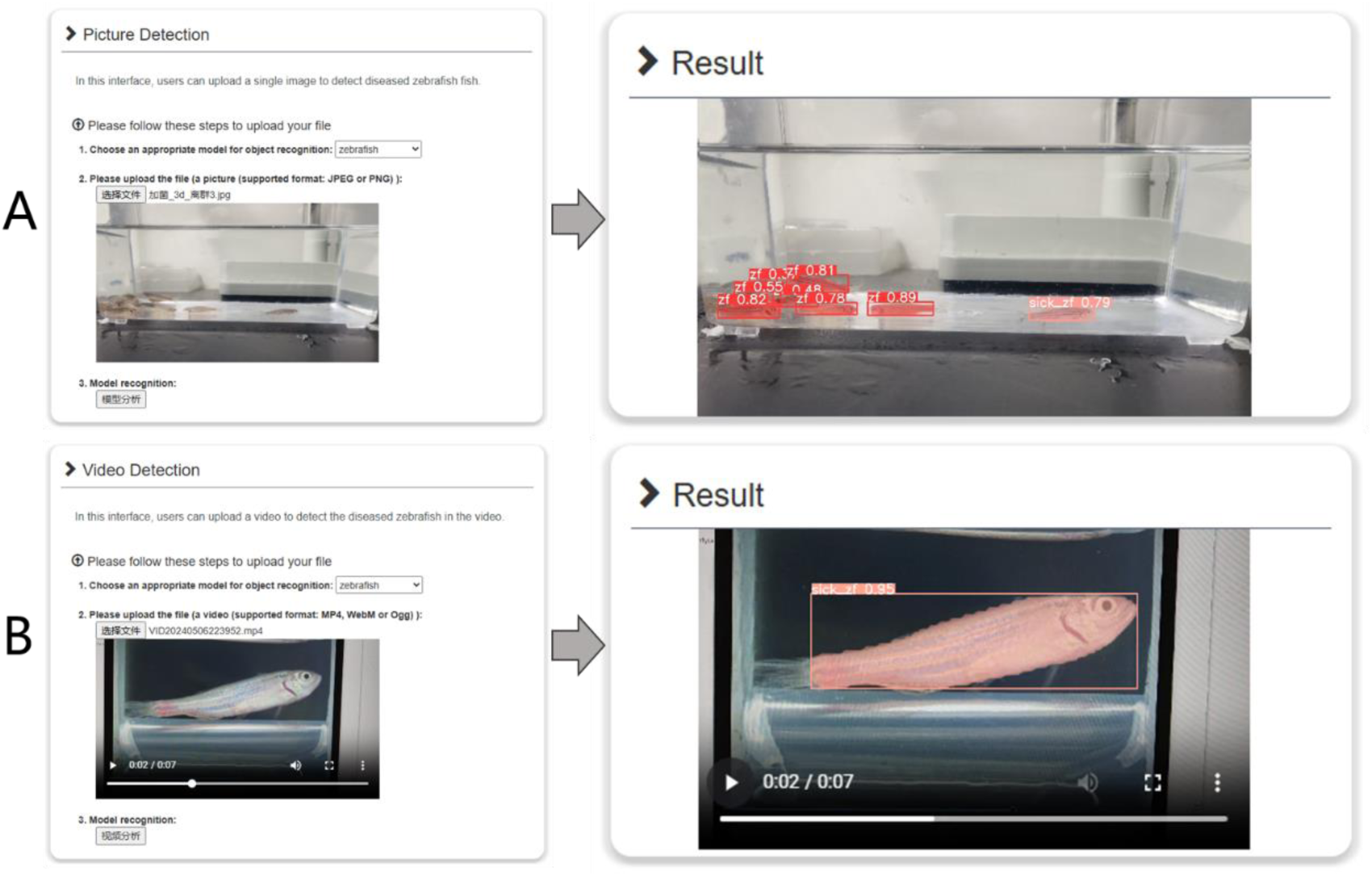
The online platform’s interface for the identification of zebrafish body surface abnormalities. A. The interface for images. B. The interface for videos.

## 4. Discussion

In this study, we addressed the challenges of detecting small targets with minimal inter-class variability and complex backgrounds in zebrafish aquaculture systems. To this end, we proposed and validated an optimized model based on YOLOv8-seg, which we termed ESC-YOLOv8-seg. Although YOLOv8-seg has already been shown to outperform other state-of-the-art models (Table 2), our method further improves detection accuracy (Table 3). By employing data augmentation and image preprocessing techniques, we have successfully mitigated the challenges posed by the subtle characteristics of zebrafish surface abnormalities and their susceptibility to environmental interference, thereby enhancing the model’s robustness.

Specifically, we conducted a comparative analysis of three advanced instance segmentation models. In comparison to Mask R-CNN and YOLACT, YOLOv8-seg demonstrated superior detection accuracy and speed, particularly in terms of detection speed, achieving up to 179 FPS, whereas Mask R-CNN only reached 5 FPS. This discrepancy can be attributed to the fact that Mask R-CNN is a two-stage instance segmentation model with a more complex overall framework and detection logic, leading to slower detection speeds. In contrast, YOLOv8-seg and YOLACT, as single-stage instance segmentation models, achieve higher frame rates through their efficient architectural design and parallel processing mechanisms, making them suitable for real-time detection scenarios. Additionally, the YOLOv8-seg model has a weight of just 6.45 MB. A smaller model weight results in fewer parameters, faster inference, and lower latency, which is crucial for applications requiring immediate responses. Furthermore, it has reduced hardware requirements, facilitating easier deployment. Consequently, YOLOv8-seg was selected as the base architecture for our improvements. As shown in Table 2 and Figure 6, the performance of YOLOv8-seg model in detecting the body surface conditions of zebrafish in real environments remains inadequate. The mAP@0.5:0.95 values for object detection (box) and instance segmentation (mask) are 0.853 and 0.702, respectively, indicating a significant gap in the model’s ability to accurately identify and segment zebrafish body surfaces in natural settings. One reason for the relatively low mAP values is that the distinguishing features between normal and abnormal zebrafish body surfaces are not as pronounced when detecting smaller targets. Abnormalities often manifest as subtle changes in body surface coloration, which can be difficult to discern. The original YOLOv8-seg model did not incorporate enhancements to the positional information of zebrafish or local surface information during the detection process. Therefore, we integrated the EMA attention mechanism into the network. The incorporation of the attention mechanism increased the number of convolutional layers, which, in the context of object detection, introduces a heightened level of difficulty in identifying smaller targets [43]. As the depth of convolution increases, the features of larger objects are more readily retained, while those of smaller objects are more easily overlooked [44]. To address the potential issues associated with dereliction detection in small targets, such as zebrafish, and those occurring when fish are distant from the camera, we implemented a modification to the SPPF module at the base of the backbone by replacing it with the SPPELAN module. The SPPELAN module combines the SPP technique with a local attention mechanism, enabling the capture of more detailed target features across various scales. Additionally, a dedicated detection head, designated as P2, was incorporated into the network to enhance the identification of smaller targets. The incorporation of the C2f_Faster module effectively mitigates the computational slowdown resulting from the increased number of convolutional layers due to the EMA attention mechanism. Despite the elevated complexity of the modified model, the single-stage parallel detection design of YOLOv8 ensures that its FPS remain sufficient for real-time detection.

The preprocessing of images is a relatively time-consuming process. The enhancement in detection accuracy and recall from the raw schooling-fish images to the preprocessed schooling-fish images dataset is comparatively limited (Figure 7). This may be attributed to the high-clarity images used in our training set, which were extracted from video sequences, thus limiting the potential for further quality enhancement through preprocessing. In practical applications, it is expected that the benefits of preprocessing will be more pronounced. While the preprocessing of training data is a one-time task that does not impact the system’s performance in real-world use, the decision to preprocess real-time data should consider the shooting conditions of the actual aquaculture environment and computational hardware capabilities.

Although our study has made significant progress in accurately detecting small targets in real time, several challenges remain for practical applications. For example, the majority of the diseased zebrafish image data was obtained from experiments involving *Aeromonas hydrophila*, with a smaller portion derived from disease outbreaks in fish labs. The diversity of surface abnormality types is somewhat limited, and the model’s generalization capability still requires further validation. Additionally, in zebrafish farming facilities, numerous aquariums that require monitoring, which places higher demands on both software and hardware. Furthermore, with aquariums closely positioned, determining the optimal angle for capturing clearer image data presents a new challenge. To enhance practicality, the following improvements and expansions can be considered:

1. Dataset Expansion: Collect a more diverse dataset of zebrafish surface abnormalities caused by various etiologies (infectious and non-infectious) under different scenarios, such as varying lighting conditions, stocking densities, and fish ages. This will enrich the training dataset and improve the model’s generalization capabilities.
2. Model Optimization: Further refine the model architecture to reduce computational complexity and enhance real-time detection performance. Additionally, explore the integration of advanced techniques, such as incorporating Transformers [45], variants of Transformers [46], and other attention mechanisms.
3. Multimodal Data Fusion [47]: Integrate other types of data, such as water quality, behavioral trajectories, and biomass estimations, to develop a multimodal anomaly detection system that improves the accuracy and reliability of detections.
4. Cross-species Generalization: Validate and adapt the model for the surface anomaly detection of other economically important fish species, thereby expanding its scope of application.

Additionally, we have conducted a preliminarily exploration of the model’s potential for identifying diseased regions in zebrafish. Despite the basic classification of the diseased areas, the results were promising, with an accuracy rate of 91.44%. However, it is important to note that YOLOv8-cls is designed for whole-image classification and can only categorize the entire image, identifying the most probable class for the entire image. In instances where multiple lesion areas are present within a single image, as illustrated by the red boxes in Figure 11, the classification may not be entirely accurate. With a more diverse range of disease types and corresponding image data, the model has the potential to be further refined for the classification of disease types and for more precise detection of diseased fish, which could significantly enhance the regulation and improvement of fish welfare.

The methodology developed in this study, while specifically designed for the health monitoring of zebrafish, holds promise for adaptation to the real-time surveillance of economically important fish species in industrial aquaculture settings. By employing targeted image acquisition devices and adjusting model parameters, the system can facilitate the early detection and timely management of surface abnormalities in economic fish species. Early identification and prompt intervention can significantly reduce the risk of disease transmission and ensure the healthy development of aquaculture industries.

## 5. Conclusions

To address the challenges associated with the monitoring and early warning of surface abnormalities in intensive recirculating aquaculture systems for zebrafish, this study proposes a computer vision-based real-time detection method, ESC-YOLOv8-seg. This method enables the real-time monitoring of surface abnormalities in zebrafish within recirculating aquaculture systems, providing a robust health safeguard scheme for zebrafish, which are increasingly utilized in research. The incorporation of image preprocessing, high-resolution images, and model modifications enables ESC-YOLOv8-seg to detect subtle differences on small targets. Experimental results demonstrate that the model effectively identifies abnormalities on the surface of zebrafish and exhibits high detection accuracy and stability across various test datasets. This makes it suitable for assessing fish surface conditions caused by diseases, water quality changes, and other factors. In future research, the methodology presented in this study may be extended to the industrial farming of economically important fish species through the expansion of datasets and fine-tuning of the model.

## Ethics statement

All experiments and animal treatments were carried out according to the principles of Animal Care and Use Committee of Institute of Hydrobiology, Chinese Academy of Sciences.

## CRediT authorship contribution statement

**Danying Cao**: Conceptualization, Methodology, Software, Formal analysis, Investigation, Writing - original draft. **Hong Yang**: Methodology, Data curation. **Cheng Guo**: Software, Data curation. **Yingyin Cheng**: Validation, Resources, Supervision. **Wanting Zhang**: Data curation. **Mijuan Shi**: Conceptualization, Investigation, Writing - review & editing. **Xiao-Qin Xia**: Conceptualization, Investigation, Writing - review & editing, Project administration, Funding acquisition.

## Acknowledgments

This work was supported by the National Key R&D Program of China (Grant No. 2021YFD1200804), the National Key R&D Program of China (Grant No. 2023YFD2401603), the Key R&D Program of Shandong Province, China (Grant No. 2023LZGC020), and the Foundation of State Key Laboratory of Mariculture Biobreeding and Sustainable Goods (Grant No. BRESG202306).

## Data Availability

Data will be made available on request.

