## Supplementary Figures for "A real-time detection and non-destructive warning method for zebrafish body surface anomalies based on improved YOLOv8 framework"

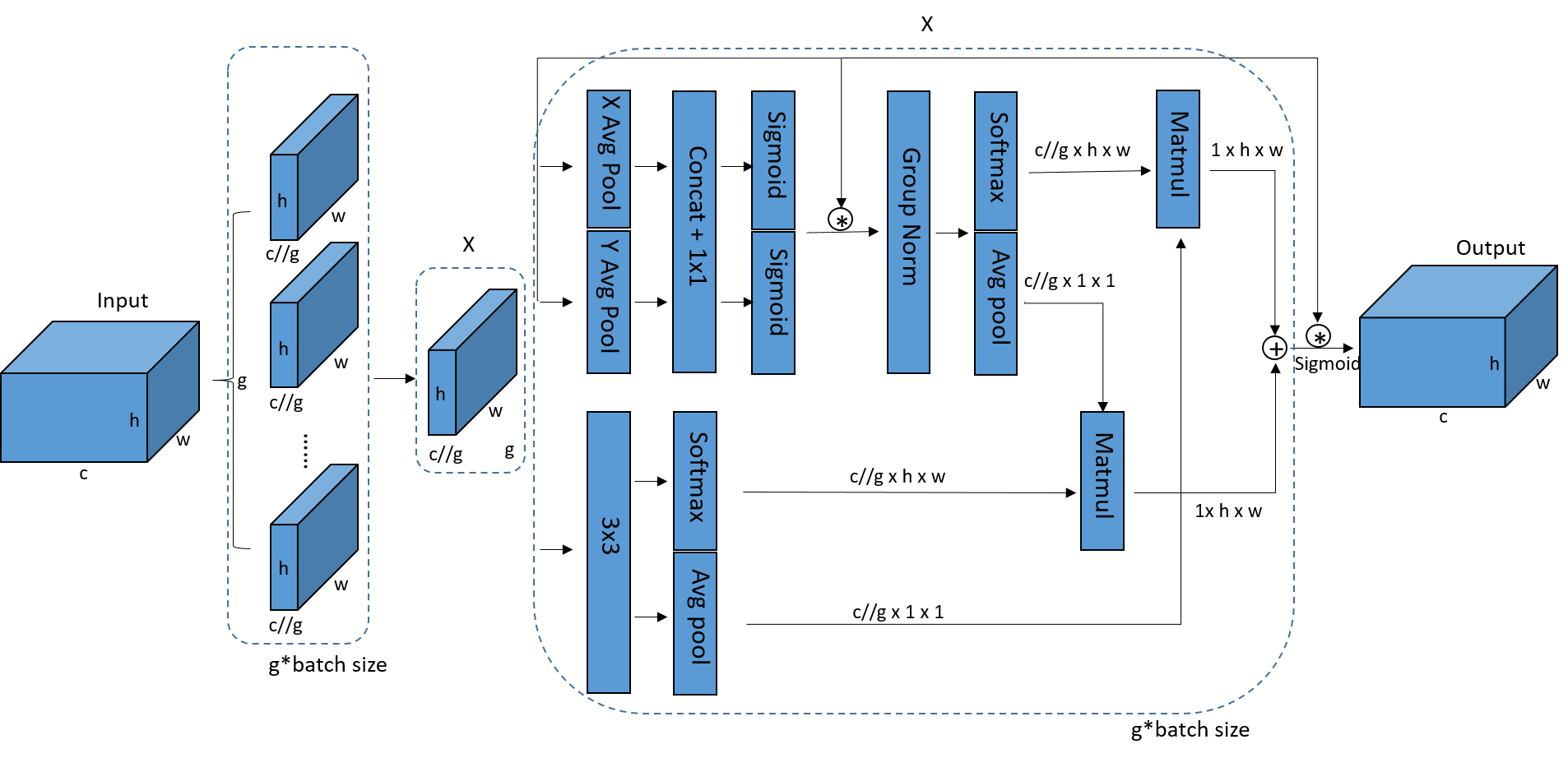

Supplementary Figures 1. Illustration of the EMA. Here, “g” means the divided groups, “X Avg Pool” represents the 1D horizontal global pooling and “Y Avg Pool” indicates the 1D vertical global pooling, respectively.

The EMA module utilizes a parallel substructure. For any given input feature map, EMA divides it along the channel dimension into g sub-features, where g is significantly smaller than the total number of channels c. The learned attention weight descriptors are used to enhance the feature representation within each sub-feature. For each grouped feature map g, EMA extracts its attention weight descriptor through three parallel pathways. Two of these parallel pathways are in the 1×1 branch, where the structure improves upon the sequential processing method of CA. The third pathway is located in the 3×3 branch, where the larger local receptive fields facilitate the capture of multi-scale spatial information. In the 1×1 branch, two 1D global average pooling operations are applied to encod the channel along two spatial dimensions, while in the 3×3 branch, a single 3×3 convolutional kernel is stacked to capture multi-scale feature representations. After obtaining the feature representation of the region of interest, EMA introduces two tensors to aggregate cross-spatial information from different spatial dimensions. One tensor is the derived from the 1×1 branch, and the other is from the 3×3 branch. The global spatial information is encoded in the 1×1 branch using 2D global average pooling, while in the 3×3 branch, the global spatial information is similarly encoded. Finally, the output of EMA has the same size as the input.
